# Direct Observation of Topoisomerase IA Gate Dynamics

**DOI:** 10.1101/356659

**Authors:** Maria Mills, Yuk-Ching Tse-Dinh, Keir C. Neuman

## Abstract

Type IA topoisomerases cleave single-stranded DNA and relieve negative supercoils in discrete steps corresponding to the passage of the intact DNA strand through the cleaved strand. Although it is assumed type IA topoisomerases accomplish this strand passage via a protein-mediated DNA gate, opening of this gate has never been observed. We developed a single-molecule assay to directly measure gate opening of the *E. coli* type IA topoisomerases I and III. We found that following cleavage of single-stranded DNA, the protein gate opens by as much as 6.6 nm and can close against forces in excess of 16 pN. Key differences in the cleavage, ligation and gate dynamics of these two enzymes provide insights into their different cellular functions. The single-molecule results are broadly consistent with conformational changes obtained from molecular dynamics simulations. These results allow us to develop a mechanistic model of type IA topoisomerase-ssDNA interactions.

## Introduction

DNA topology in the cell is controlled by a group of DNA-remodeling enzymes known as topoisomerases^1,2^. Topoisomerases are divided into two classes: type II topoisomerases cleave both strands of duplex DNA (dsDNA) and pass a second duplex through the transient break^3^, whereas type I topoisomerases cleave single-stranded DNA (ssDNA) and either permit rotation of the intact strand (Type IB)^4,5^ or pass the intact strand through the transient nick (Type IA)^6,7^. Type IA topoisomerases are ubiquitous and essential enzymes that include the bacterial topoisomerases I^8^ and III^9^, the archaeal reverse gyrase^10^, and the eukaryotic topoisomerases IIIα^11,12^ and IIIβ^13^ Biochemical activities of type IA topoisomerases include relaxation of negatively supercoiled DNA^14^, decatenation of linked DNA^15,16^, and the addition of knots^17^. In prokaryotes type IA topoisomerases are involved in relaxation of excess negative supercoils^18^ and resolution of replication and recombination intermediates^19,20^. In eukaryotes, these enzymes are involved in accurate chromosome segregation during mitosis^12,21,22^, non-crossover resolution of double Holliday junctions^23,24^ and interactions with mRNA to promote neurodevelopment and prevent mental dysfunction^25-27^.

Type IA topoisomerases bind and cleave single-stranded DNA and then pass a second single strand or duplex through the break, changing the linking number by 1^7^. This strand passage mechanism has long been assumed to involve a protein-mediated DNA gate^28-31^. In the gate model, ssDNA is bound across domains I and IV of the topoisomerase and is cleaved by the active site tyrosine in domain III (**Fig. 1a-b**)^32^. Domain III then moves away from domain I, creating an opening through which a second strand of DNA can pass. The enzyme must then return to the closed state, religate the DNA backbone, and open again to release the trapped DNA (**Fig. 1b**). Although structural,^31,32^ biochemical,^6,33^ and single-molecule experiments^7^ support this model, direct evidence for the gate mechanism has proved elusive. No structures exist of a type IA topoisomerase in the open state and no measurements have definitively shown that an open state exists, leaving open the possibility that the strand passage is coupled to some other conformational change^34^.

**Figure 1.**
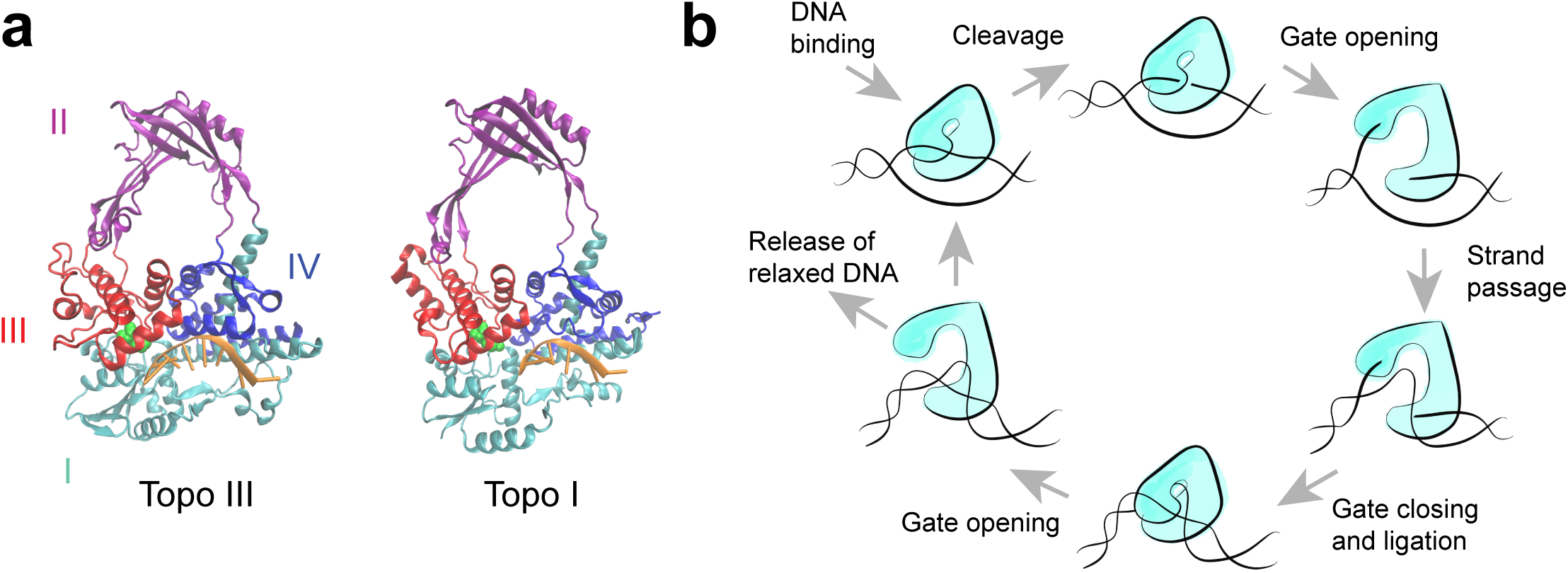
Proposed mechanism of type IA topoisomerase activity. **(a)** Crystal structures of the *E. coli* type IA topoisomerases, topo III (pdbid: I17D)^49^ and topo I (pdbid: 1MW8)^37^. The domains are labeled according to color. DNA bound to the active site is shown in orange and the catalytic tyrosines are highlighted in green. For simplicity, the C-terminal zinc finger domains of topo I are not shown. (**b**) Proposed model of type IA topoisomerase protein-mediated DNA gate mechanism for strand passage.

Here we describe the direct observation of type IA topoisomerase gate opening dynamics in single-molecule magnetic tweezers based measurements of the two *E. coli* type IA topoisomerases, topo I and topo III, interacting with ssDNA (**Fig. 2a** and **3a**). By analyzing the force- and magnesium-dependent kinetics of the gate opening and closing, we delineate the catalytic cycle of type IA topoisomerases. These results show key differences in the gate dynamics of topo I and topo III that may underlie the differences in their biochemical activities. Molecular dynamics simulations reveal the potential structure of the open conformation and confirm that the experimentally observed extension changes are consistent with opening of the gate between domains I and III. Our results provide a detailed description of the kinetics of type IA topoisomerase-ssDNA interactions and establish an experimental paradigm to study the conformational dynamics of type IA topoisomerases.

**Figure 2.**
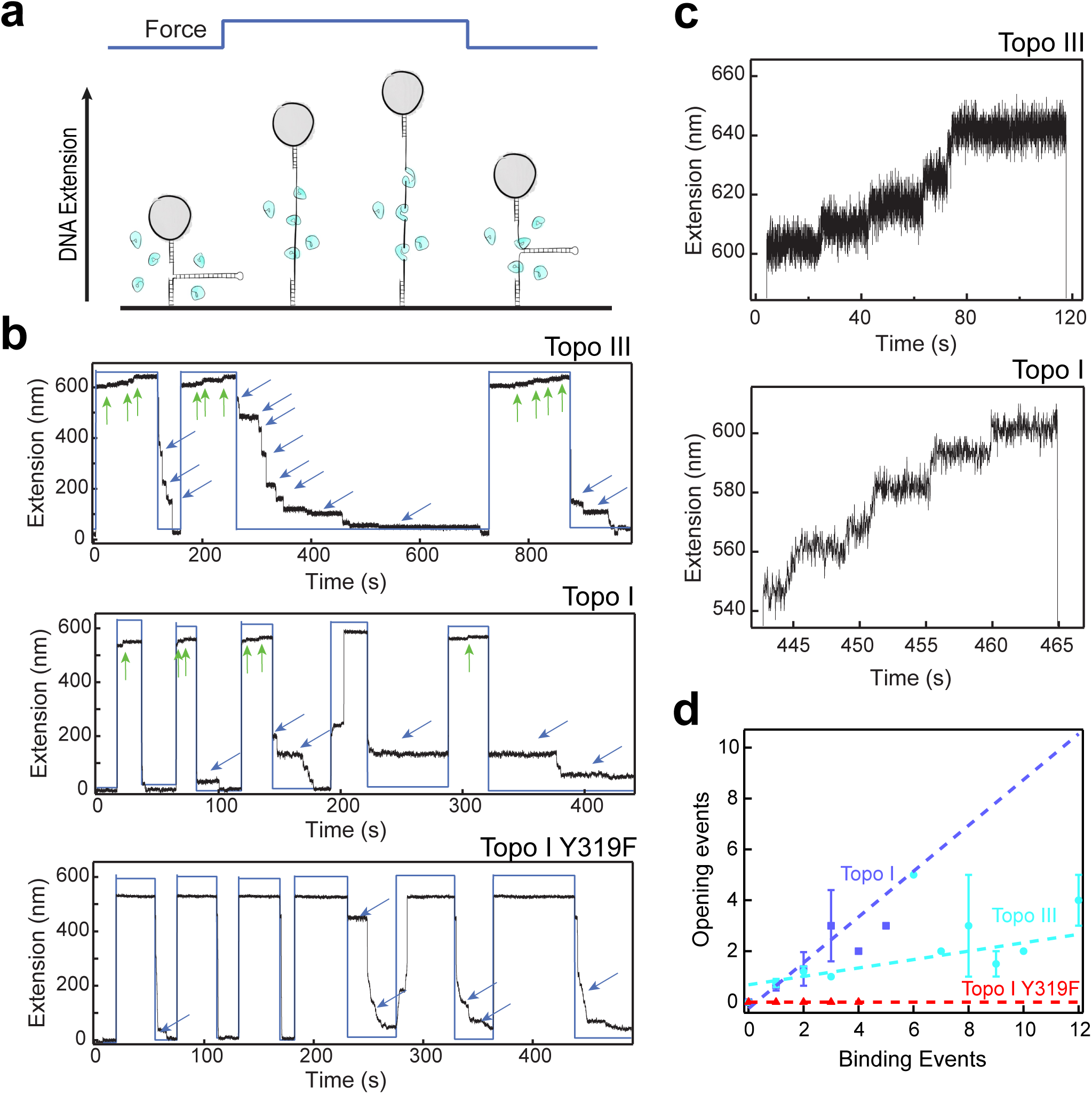
Magnetic tweezers measurements of topoisomerase binding and gate opening. (**a**) Experimental design. Hairpin DNA was unfolded at high force in the presence of topoisomerase. Gate opening was detected as an increase in DNA extension beyond the fully extended state. Binding was detected as pauses in refolding upon lowering the force. (**b**) Example experimental traces for topo III, topo I, and topo I Y319F. Black lines are DNA extension. Blue lines represent the force, alternating between ∼9 pN and 22 pN. Green arrows indicate gate opening events. Blue arrows indicate binding. Differences in the baseline extension of the unfolded state are due to variances in the applied force. (**c**) Expanded examples of gate opening events for topo III and topo I. Extension increases were approximately 6.6 ± 0.7 nm for topo III and 6.6 ± 1.0 nm for topo I (**Supplementary Fig. 1**). (**d**) Correlation between gate opening and topoisomerase binding events. For each opening and closing transition, the number of opening and binding events were scored. The average number of opening events for a given number of binding events is plotted. Topo III is shown in cyan (*n_tethers_* = 4, *n_events_* = 58), topo I in dark blue (*n_tethers_* = 5, *n_events_* = 25), and topo I catalytic mutant Y319F in red (*n_tethers_* = 5, *n_events_* = 18). Error bars are standard deviations. Dashed lines indicate linear fits to the data with slope 0.16 ± 0.05 for topo III, 0.9 ± 0.5 for topo I, and 0.0 for topo I Y319F.

**Figure 3.**
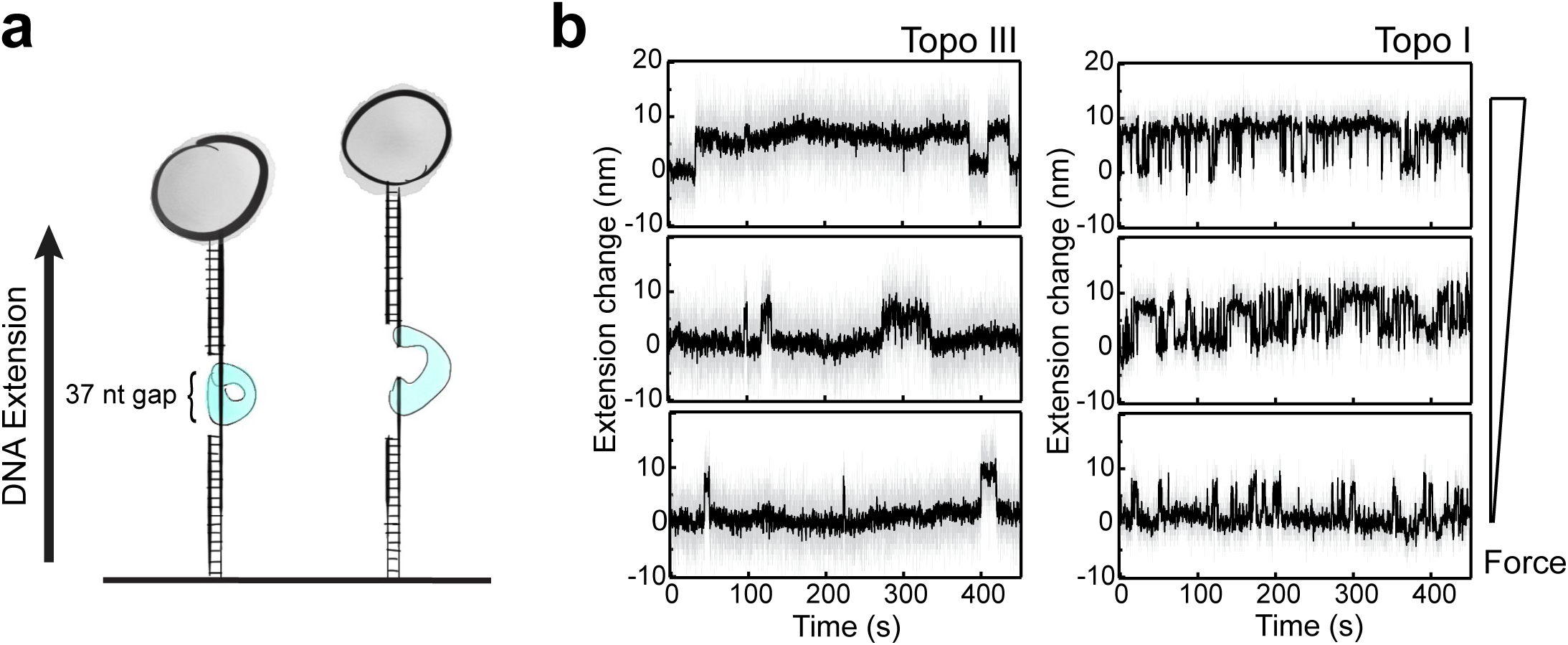
Force dependence of gate opening dynamics. (**a**) Experimental design. Duplex DNA with a 37 nt gap was held at constant force and gate opening/closing detected as transient increases in extension of ∼ 6 nm. (**b**) Example traces showing extension change of single DNA molecules as a function of time for topo III and topo I at 16, 14, and 12 pN. Gray lines are raw data. Black lines are smoothed data using a 4^th^ order Savitzky-Golay algorithm over 51 points.

## Results

### Cleavage by type IA topoisomerases increases ssDNA extension by greater than 6 nm

We first tested the ssDNA binding properties of type IA topoisomerases using a single molecule magnetic tweezers^35^ hairpin rezipping assay (**Fig. 2a**)^36^. In this assay, a 537 bp hairpin was attached via dsDNA handles to a coverslip on one end and a magnetic bead on the other. The hairpin was mechanically unfolded by applying high force (∼22- 24 pN) in the presence of 500 pM *E. coli* topo III or topo I (**Fig. 2b**). The DNA was held at high force for 30-120 s after which the hairpin was allowed to refold by reducing the force (∼9 pN). Binding of topoisomerase to the DNA was detected as pauses in hairpin refolding (**Fig. 2b**, blue arrows). Subsequent dissociation of the topoisomerase resulted in further refolding of the hairpin. Pauses in refolding of the hairpin occurred at multiple distinct locations, indicating stable binding of multiple proteins.

At high force, in the presence of topoisomerases we observed an increase in the bead height beyond the extension of the fully unfolded hairpin (**Fig. 2b**, green arrows). These extension increases were on average 6.6 ± 0.7 nm for topo III and 6.6 ± 1.0 nm for topo I (**Fig. 2c, Supplementary Fig. 1**). Since the hairpin was fully unfolded at this force, we interpreted these increases in DNA extension as cleavage of the DNA by the topoisomerase followed by a protein conformational change that separated the cleaved ends of the DNA. To verify that the extension increases were cleavage-dependent, we repeated the hairpin experiments with a cleavage deficient mutant of topo I, Y319F (**Fig. 2b**)^37^. We observed refolding pauses characteristic of protein binding (**blue arrows**), but no discrete increases in extension at high force.

Comparing the number of ∼6.6 nm extension increases (opening) with the number of distinct binding events for each unfolding-refolding cycle indicates that they are correlated (**Fig. 2d**). For both topo III and topo I, the relationship between the number of extension increases and the number of bound proteins is roughly linear, with slopes of 0.16 ± 0.05 (*n_tethers_* = 4, *n_events_* = 58), and 0.9 ± 0.5 (*n_tethers_* = 5, *n_events_* = 25), respectively. The fits indicate that nearly all the bound topo I molecules cleaved and opened the DNA, whereas only a small fraction of topo III molecules did. We observed no opening events in the absence of binding (*n_tethers_* = 5, *n_events_* = 18). Based on these results, we concluded that the 6.6 nm extension increases correspond the opening of a topoisomerase-mediated DNA gate.

### Force dependence of gate dynamics

The difference in the number of cleavage events relative to binding events for the two topoisomerases in the hairpin experiments indicates differences in gate dynamics that may contribute to their different biochemical activities and distinct cellular roles. To accurately characterize the conformational changes and kinetics of the gate dynamics, we developed an assay that allowed us to measure gate dynamics of a single topoisomerase bound to ssDNA as function of force (**Fig. 3a**). Force selectively affects the kinetic transitions involving motion by altering the underlying free energy profile. Measuring the gate dynamics as a function of applied force therefore reveals kinetic steps involving motion and provides a measure of the free energy profile associated with the motion. For these experiments, a 2.5 kb dsDNA with a 37 nt gap was attached to a coverslip on one end and a magnetic bead on the other. The limited size of the ssDNA region increases the probability that changes in the extension of the DNA correspond to the activity of a single topoisomerase.

Opening and closing of the topoisomerase gate was observed as transient increases in the DNA extension (**Fig. 3b**), *n*_tethers_ = 8. The average size of this extension change was 5.5 ± 0.4 nm for topo III, independent of the applied force (8 to 16 pN) (**Supplementary Table 1**). For topo I, the extension change increased slightly with applied force (12 to 18 pN, *n*_tethers_ = 7) (**Supplementary Table 1**), with an average value of 5.9 ± 0.6 nm. Surprisingly, the topoisomerases were able to close the gate and religate DNA against forces of 18 pN.

Whereas the overall gate dynamics are similar for both topo I and topo III, there are significant kinetic differences. To obtain kinetic states from the extension data, we used vbFRET^38^ (**Fig. 4a-b**), which applies an unbounded hidden Markov model to determine the number of states in single-molecule time traces. The program reliably found two extension states for both topo III and topo I. However, analysis of the lifetime distributions revealed a third state; in addition to the open state and a long-lived closed state, there is a short-lived closed state (**Fig. 4a-b**).

**Figure 4.**
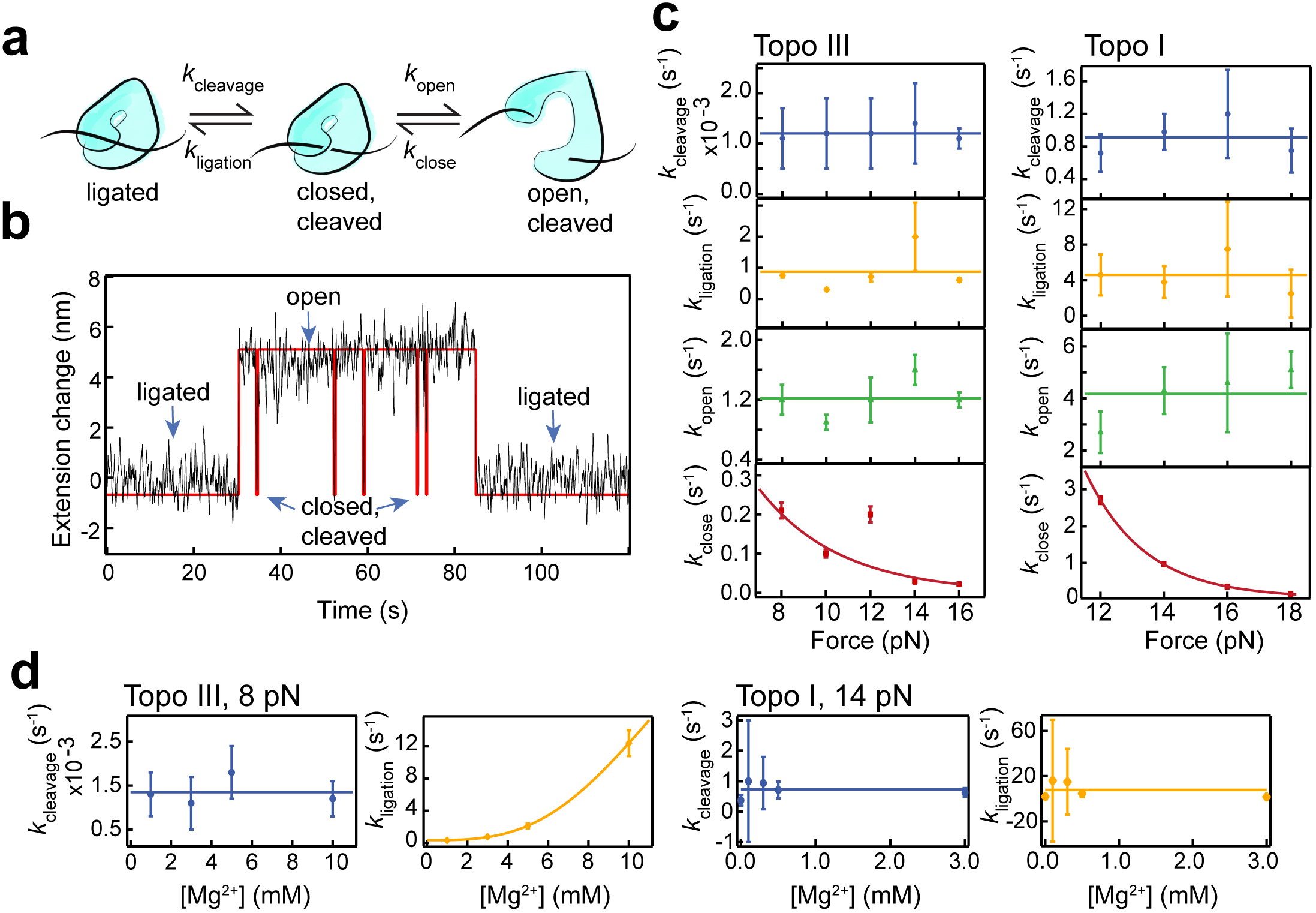
Three-state kinetics of topoisomerase IA gate dynamics. (**a**) Three-state kinetic model. (**b**) Example of a single kinetic cycle of topo III from cleavage to transient opening and closing of the gate in the cleaved state, followed by religation. Black line is smoothed data, red line is the HMM fit from vbFRET. (**c**) Force dependent kinetic analysis of *k*_cleavage_ (blue), *k*_ligation_ (orange), *k*_open_ (green), and *k*_close_ (red). Values were calculated from exponential fits of the lifetimes of each state (**Supplementary Fig. 3-4**). Error bars represent standard deviations of the fit coefficients. For *k*_cleavage_, *k*_ligation_, and *k*_open_, lines represent average values. *k*_close_ values were fit with barrier crossing model in which the force biases the kinetics of the gate closure: *k* = *k*_0_exp(-FΔx/*k*_B_T), where *k*_0_ is the force-independent closing rate, *k_B_T* is the thermal energy, and Δx is the distance between the open state well and the transition state energy barrier along the direction of applied force. The fit returned Δx and *k*_0_ values of 1.10 ± 0.04 nm and 1.8 ± 0.3 s^−1^ respectively, for topo III and 2.2 ± 0.1 nm and 1642 ± 549 s^−1^ for topo I. (**d**) Magnesium concentration dependence of ligation and cleavage rates for topo III at 8 pN force and topo I at 12 pN force. The topo III magnesium-dependent ligation rate was fit with a sigmoid with a K_m_ of 8.3 ± 1.2 mM Mg^2+^ and a V_max_ of 16.7 ± 4.7 s^−1^.

These transient closed states are consistent with a three-state model in which the protein-ssDNA complex is in a ligated state (L), a closed cleaved state (C), or an open cleaved state (O) (**Fig. 4b**), resulting in the following kinetic scheme;

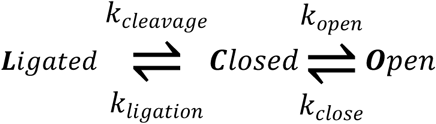

In this model, the short-lived closed states correspond to state C, which the protein may visit multiple times before religating DNA and returning to the longer-lived closed state L. Although we have assigned three states to our data, it is likely that there are additional states that we cannot directly observe from the extension measurements alone.

The gate kinetics as a function of force for both topo III and topo I are plotted in **Fig. 4c**. For topo III, the lifetimes of the long- and short-lived closed states are well-separated and the individual lifetime distributions were fit to single exponentials to determine *k*_open_ and *k*_cleavage_ (**Supplementary Fig. 3a**). For topo I, the short and long closed-state lifetimes were comparable and the closed state lifetime distribution was well-fit with a double exponential (**Supplementary Fig. 4a**). We assigned the faster rate to *k*_open_ and the slower rate to 1/(*t*_cleavage_ + *t*_open_), from which we can determine *k*_cleavage_. The distribution of open state lifetimes was fit to an exponential to determine *k*_close_ (**Supplementary Fig. 3b, 4b**).

To estimate the ligation rate, *k*_ligation_, we consider the kinetic competition between ligation and opening from the closed state C. The probability of opening from state C depends on the rates of these two competing pathways:

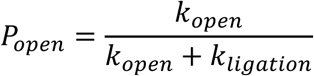

We determine P_open_ from the number of opening events (short-lived closed states) relative to the total number of closed state events.

For both topoisomerases, *k*_cleavage_ and *k*_ligation_ were insensitive to the applied force (**Fig. 4c, Table 2**). Surprisingly, the opening rates were also unaffected by the applied force (**Fig. 4c, Table 2**). This lack of force dependence indicates that the opening rate reflects a rate-limiting conformational change that precedes the mechanical opening of the gate. The closing rates exhibited an exponential force dependence (**Fig. 4c**), *k*(F) ≈ *k*_0_exp(-FΔx/*k*_B_T), where *k*_0_ is the zero-load closing rate, Δx is the distance from the open state to the transition state, and *k*_B_*T* is the thermal energy. The distances to the transition state, Δx_close_, obtained from exponential fits are 1.10 ± 0.01 nm for topo III and 2.2 ± 0.1 nm for topo I. The zero-load closing rates are 1.8 ± 0.3 s^−1^ for topo III and 1642 ± 549 s^−1^ for topo I (**Table 1**).

**Table 1.**
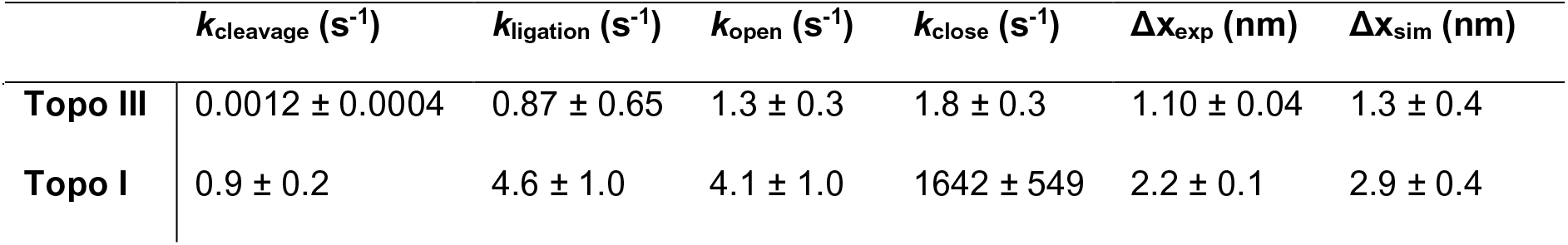
Equilibrium rates and barriers associated with Figure 6. Zero-load values for *k*_cleavage_, k_ligation_, and the rate limiting conformation changes associated with gate opening *k*_open_, and closing, *k*_close_. Δx_close_ values calculated from experiments and simulations are also shown. In the case of Topo III, this corresponds to Δx_close**_, the distance between the fully open state and the barrier associated with formation of an open-state stabilizing salt bridge.

The lifetime of the open state is several orders of magnitude longer for topo III than topo I (**Fig. 4c, Table 1**). Biochemical and single-molecule studies demonstrate that topo I is generally more effective at relaxing negatively supercoiled DNA than topo III^39^, whereas topo III is more effective at unlinking DNA catenanes^15,40^ Relaxation of negative supercoils requires passage of a single DNA strand through the protein-mediated DNA gate. In contrast, decatenation activity requires passage of dsDNA. The more stable open state of topo III could facilitate capture of duplex DNA required for decatenation, whereas the faster dynamics of topo I are compatible with capturing a local ssDNA segment. The fast closing rate also explains the difficulty in capturing the open conformation of topo I; only by slowing gate closing through the application of significant opposing forces could the open state configuration be observed.

The estimated rate of a full catalytic cycle based on extrapolated zero force kinetics of the individual steps for topo I is 0.64 ± 0.38 s^−1^, which is comparable to previous relaxation rate measurements (**Table 1, Supplementary Table 4**). Conversely the estimated rate of the topo III catalytic cycle is 0.0011 ± 0.0004 s^−1^. This unexpectedly slow rate, which is dominated by the cleavage rate, is much lower than measured relaxation rates. The slow cleavage rate is consistent with previously observed long lag times (up to ∼129 s for negatively supercoiled DNA) between relaxation events for topo III^39,40^. However, these experiments also showed higher processivity and unlinking rates for topo III during processive relaxation (between ∼19 s^−1^ and 129 s^−1^, **Table 1, Supplementary Table 4**). One possible explanation for the discrepancy is the absence of a second DNA strand in our experiments. If topo III cleavage and ligation rates were stimulated by the presence of a second ss- or ds-DNA strand, then passage of an initial strand could lead to the fast bursts of processive activity previously observed^39^. The requirement of a second strand of DNA would align with kinetic studies of the S. *solfatarius* topo III in which ligation of cleaved ssDNA was facilitated by annealing of a complementary strand^41^.

### Magnesium dependent kinetics

Although magnesium is necessary for the activity of type IA topoisomerases, the distinct roles that magnesium plays in cleavage and religation remain under debate. Previous studies have shown that increasing magnesium concentration increases the cleavage product, but magnesium is not required for cleavage^42^. Religation requires magnesium, but the magnesium dependence of the religation rate is unknown ^43,44^. Ensemble measurements report changes in steady state levels of cleaved and religated product, but do not provide direct information on the cleavage or ligation rates. Stopped flow experiments offer a more direct way to measure these kinetics, but to date few such studies have been done on type IA topoisomerases^41^. We determined the effect of magnesium on the cleavage and religation rates by measuring the gate kinetics as a function of magnesium concentration (**Fig 4d, Table 3**).

The cleavage rate of both proteins was independent of the magnesium concentration (**Fig 4d, Supplementary Table 3**). The ligation rate of topo III depended strongly on the magnesium concentration (**Fig. 4d, Supplementary Table 3**), exhibiting a sigmoidal relationship, with a K_m_ of 8.3 ± 1.2 mM Mg^2+^ and a V_max_ of 16.7 ± 4.7 s^−1^. This is close to previously measured values of the K_m_ for the magnesium dependence of relaxation by type IA enzymes, which range from 2-4 mM^9,41,45^. The fact that the relationship is sigmoidal rather than hyperbolic is consistent with biochemical evidence indicating that the catalytic cycle depends on two coordinating magnesium ions^45,46^. In contrast, the ligation rate of topo I showed no magnesium dependence (**Fig. 4d, Supplementary Table 3**). The differences in magnesium dependence of religation may be due to differences in the active sites of the two topoisomerases. Topo I may have one tightly bound Mg^2+^ under these reaction conditions^42,43^. Topo III has a strictly conserved Lysine residue (K8) that interacts with the putative scissile phosphate in the active site^44^, and as a result may not bind magnesium as tightly as topo I. Based on a recently determined structure of *M. tuberculosis* topoisomerase I with bound ssDNA and Mg^2+^, it was proposed that the observed Mg^2+^ may play a role in correctly positioning the 3'-hydroxyl group relative to the phosphotyrosine linkage for DNA religation. The potential role of a second Mg^2+^ remains to be determined.

For topo III we observed a decrease in binding affinity and reduced religation below 1 mM Mg^2+^, which precluded measurements at lower magnesium concentrations. Topo I showed decreased binding below 0.3 mM, but was still able to cleave and religate DNA, even without added magnesium in the buffer. Only when EDTA was added to the reaction to chelate trace amounts of metal was ligation by topoisomerase I suppressed (**Supplementary Fig. 5**). These results suggest that magnesium-dependent DNA binding effects may contribute to observed effects of magnesium on the ratio of cleavage to religation from ensemble measurements.

### Gate dynamics from simulations

Our experiments indicate a separation between the cleaved DNA ends of 5.5 - 6.6 nm, which is unexpectedly large. Single-stranded DNA needs a gap of only ∼1 nm to pass through the break for relaxation, whereas duplex DNA needs ∼2 nm. Given the high forces used in our magnetic tweezers experiments, it is possible that the enzyme normally undergoes a much smaller conformational change in the absence of force on the DNA. To test this possibility, we conducted umbrella sampling^47,48^ molecular dynamics simulations of topo III bound to ssDNA (**Fig. 5, Supplementary Movie 1**)^49^. In these simulations, the distance between the center of mass of domain III (shown in red in **Fig. 5a**) and the center of mass of domain IV (shown in blue) was gradually increased over a series of windows. These domains must move away from each other for the gate to open, but the direction of this motion may not be aligned with our experimentally imposed force and displacement axis. Restraining the distance between the domains rather than pulling along a vector should allow the system to follow an energetically unbiased pathway.

**Figure 5.**
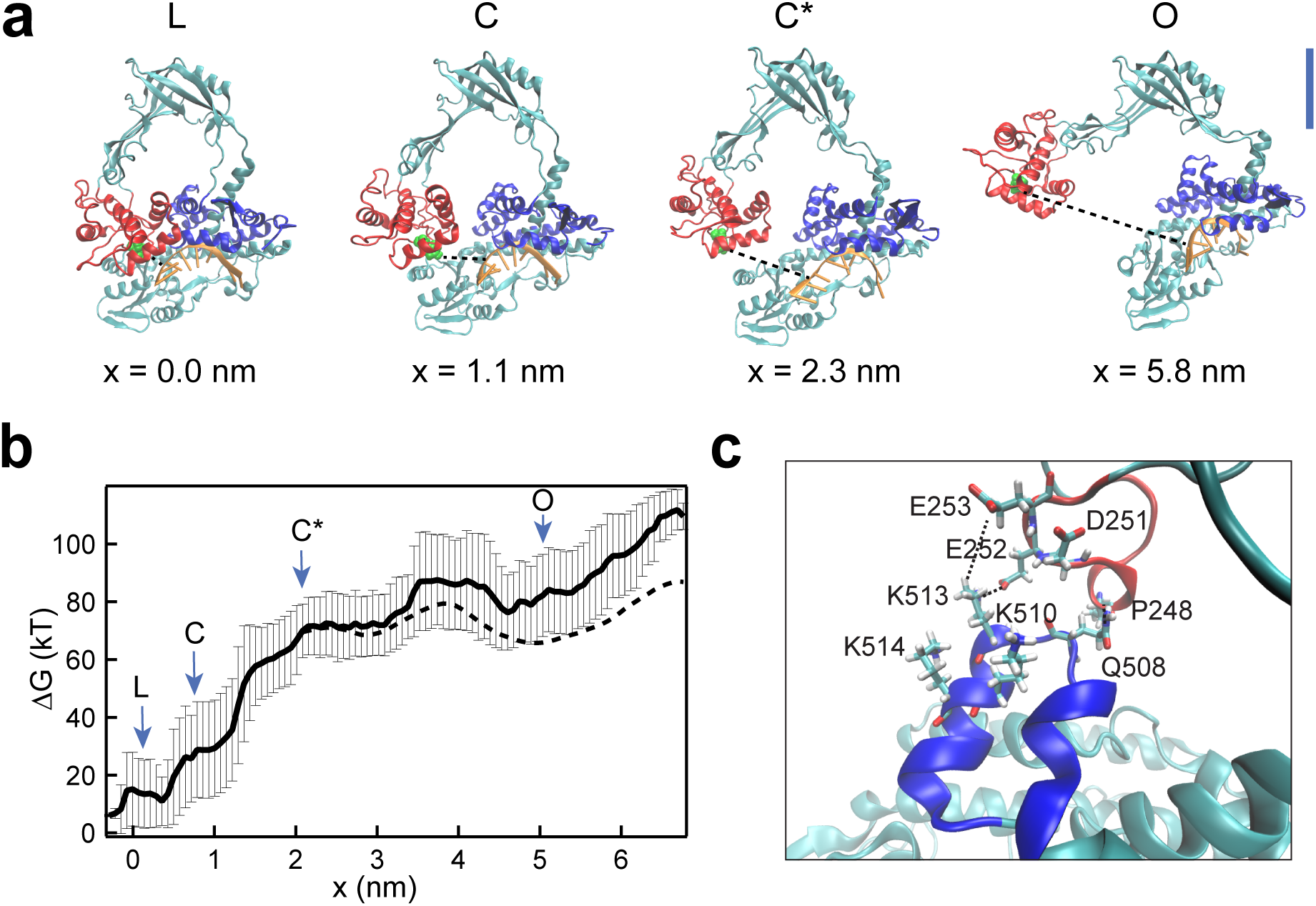
Molecular dynamics simulations of topo III gate opening. (**a**) Topo III structures from molecular dynamics umbrella sampling simulations. Representative structures are shown for the closed state (L), the state in which contacts between domains III and IV are broken (C), the state in which contacts between domains III and I are broken (C*), and the fully open structure (O). x (dashed black line) is the change in distance between the oxygen of the catalytic tyrosine, Y328, and the cleavage site of the DNA backbone. Blue scale bar represents 2 nm. (**b**) Relative free energy change as a function of gate opening (x). Arrows indicate the location of the states represented in (**a**). The dashed line is the estimated free energy for an applied force of 22 pN under the assumption that the transition from L to C* is independent of force (**Supplementary Fig. 6**). (**c**) Structure of the decatenation latch. The decatenation loop is shown in blue and the acidic loop in red. Key residues in forming the latch are highlighted. Dashed lines indicate specific contacts. K513 forms contacts with both E253 and E252, while Q508 interacts with the backbone of P248. Two other basic residues in the decatenation loop, K510 and K514, and a third acidic residue in the acidic loop, D251, are also involved in loop contacts at other points in the simulations (**Supplementary Movie 2**).

The simulations reveal that domain III initially moves along domain I rather than outward, first breaking contacts with domain IV (**Fig. 5a**, structure C) before breaking contacts with domain I (structure C*). As a result of this sliding motion, the protein gate is not large enough to accommodate ssDNA until the distance between the cleavage site and the catalytic tyrosine exceeds 3.8 nm. For dsDNA to pass through the DNA break, a separation of ∼5.8 nm was required (**Fig. 5a**, structure O). This separation is in excellent agreement with our experiments.

The umbrella sampling method can be used to calculate equilibrium free energy profiles along any reaction coordinate, provided the conformational space of the coordinate is fully sampled during the simulations^50^. In this way, we could estimate the free energy profile along the experimental reaction coordinate defined by the applied force (**Fig. 5b**). The simulations indicate that the initial barrier to opening is defined by breaking of contacts between protein domain III and domains I and IV. Once the interdomain contacts are broken, the free energy profile is relatively flat. There is a small barrier between ∼3.5 and 4.5 nm that corresponds to interactions between the decatenation loop of topo III and a second smaller loop on domain II, both of which are absent in topo I (**Fig. 5c, Supplementary Movie 2**). The decatenation loop has a cluster of conserved basic residues^51^, which form contacts with acidic residues in the second loop. The interaction presumably stabilizes the open state and suggests a structural role for the decatenation loop, which is required for the unlinking activity of topo III^51^. The stabilization of the open state explains why the closing rate is slower for topo III in our experiments (**Fig. 4c, Supplementary Table 2**). Moreover, the distance of this barrier from the open state well is 1.3 ± 0.4 nm, which is in excellent agreement with the experimental topoisomerase III Δx_close_ value of 1.1 nm. The experimental Δx_close_ for topo I, 2.2 nm, is in good agreement with the distance between the open state well and state C*, 2.9 ± 0.4 nm. The distance between the closed state and C* in the simulations, Δx_open_ is 2.2 ± 0.3 nm (**Fig. 5b**).

### Mechanistic Model of Type IA topoisomerase-ssDNA interactions

Whereas the three-state model is consistent with the number of observed states, analysis of the force-dependent rates, combined with simulation results, suggest that there is at least one additional kinetic state in the gate dynamics pathway. The experimental opening rate is independent of force (**Fig. 4C, Supplementary Table 2**). This lack of force dependence indicates that the opening rate is dominated by a rate limiting step that proceeds the mechanical opening of the gate. This additional rate limiting kinetic transition is supported by the fact that the initial barrier associated with breaking contacts between domains III and I in the simulations is too high to be thermally accessible. This high energy barrier suggests that opening is limited by a conformational change that lowers the transition energy barrier but that is too slow to be sampled in our simulations.

To accommodate these results, we extended the model of topoisomerase-ssDNA interactions (**Fig. 6, Table 1**) to include an additional state, C* that is inferred but not directly observed.

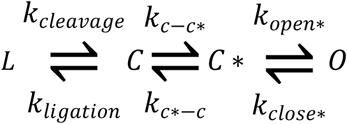

**Figure 6.**
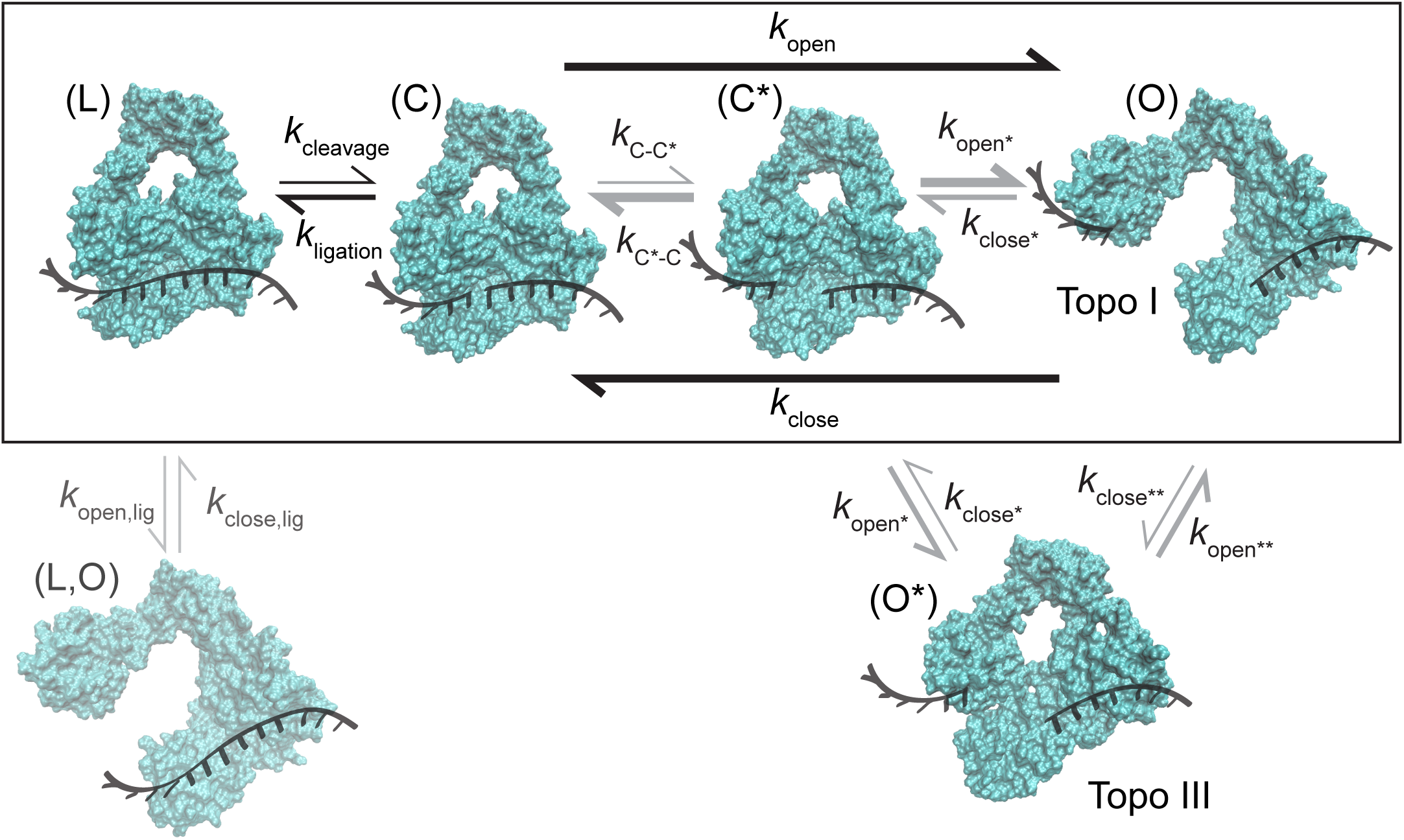
Kinetic scheme of type IA topoisomerase gate dynamics. Structures of topo III from MD simulations representing protein-ssDNA states: (L,O) open gate with intact DNA (invisible in our experiments); (L) closed gate intact (ligated) DNA; (C) closed gate with cleaved DNA; (C*) closed gate with cleaved DNA after a rate-limiting conformational change in which domain I moves relative to domain III; and (O) open gate with cleaved and separated DNA. The black box designates states common to topo I and topo III that are directly observed (L,C,O) or inferred (C*). (O*) is a topo III specific intermediate, identified from MD simulations, in which the decatenation loop forms contacts with a second loop in domain II (**Fig. 5**). Experimentally measured rates (black) are shown in **Table 4**. Rates that are inferred but not directly measured are shown in gray. The weight of the arrows represents relative kinetic rates.

Experimental cleavage and ligation rates in this scenario correspond to the chemical reactions, but the opening and closing rates each represent a convolution of two rates from C to O via C* and vice versa. Accordingly, the measured *k*_open_ is dominated by the rate of a conformational change (*k*_c-c*_) that allows domain III to escape contacts with domain I. The actual opening rate (*k*_open*_) is force dependent but too fast to observe. Similarly, *k*_c*-c_ is assumed faster than *k*_close*_ and *k*_open*_, such that the measured closing rate is dominated by *k*_close*_, and the equilibrium between C* and C is biased towards state C from which religation can occur. In this scenario, the gate can undergo multiple transitions between states C and C* before opening, in which case *k*_open_ would represent the escape rate, rather than *k*_c-c*_. The measured force dependence of the closing rate supports the hypothesis that the transition from state C* to C is fast (much faster than the fastest closing rate of 2.7 ± 0.1 s^−1^). Furthermore, the lack of force dependence of the religation and opening rates is consistent with the hypothesis that the rate *k*_c*-c_ exceeds all other rates.

We further propose that topo III has an additional state, O*, between C* and O, in which the decatenation loop forms a bridge that creates a barrier to closing. This additional barrier observed in the simulated free energy profile is supported by the significantly slower gate closing rate, and by the shorter distance between the open state and the transition state obtained from the force-dependent closing rate of topo IIII in comparison to topo I. The long lifetimes of the topo III ligated state also suggest the possibility that the protein could open and close while bound to ligated DNA (**Fig. 6**), contributing to the slower apparent cleavage rate.

## Discussion

Since the first structure of a type IA topoisomerase was solved, it has been predicted that these enzymes utilize a protein-mediated DNA gate mechanism for strand passage, similar to type II topoisomerases^28-31^. In the intervening 24 years, no structures of the protein in an open state have been solved, though crystal structures of a fragment suggest that domain II could bend, separating domains I and III^31^ (**Supplementary Fig. 7**). Biochemical studies have shown that domains I and III move relative to each other in order for strand passage to occur^6^. The same studies demonstrated that circular DNA can be trapped in a complex with the protein by crosslinking domains I and III, suggesting that the passed strand resides within the protein cavity^6^. Ensemble fluorescence quenching assays have similarly linked strand passage to a conformational change between domains I and III^33^. More recent combined magnetic tweezers - fluorescence experiments also indicate a conformational change prior to strand passage^34^, though the results were interpreted as domain closure rather than domain opening. Attempts to estimate gate opening of topo I from force-dependent relaxation kinetics resulted in a Δx value of ∼ 10 nm, too large to be reflective of gate opening^7^. These studies are consistent with, but did not conclusively demonstrate, the existence of the putative protein-mediated DNA gate or the gate dynamics. The lack of direct evidence for an open state has led to the proposal of at least one alternative model in which domains I and III move against, rather than away from, each other^34^.

By measuring the extension of individual DNA molecules containing ssDNA regions we observed conformational changes corresponding to the separation of cleaved DNA by 5.5-6.6 nm in two bacterial type IA topoisomerases. These experiments provide direct evidence for the original protein-gate model of topoisomerase IA activity. Although the magnitude of gate opening was unexpectedly large, molecular dynamics simulations indicate that a large separation is required to pass duplex DNA through the gate given the position of the DNA binding site within domain I. Whereas it is possible that the applied force resulted in larger gate separation than occurs at zero force, the fact that the gate opening and ssDNA cleavage are reversible and occur with comparable kinetics as the strand passage reaction suggest that the observed open state is biochemically relevant. This conclusion is bolstered by the close agreement between the experiments and simulations, and the fact that topo III and topo I remain catalytically active under applied loads as high as 12 pN (**Supplementary Fig. 8**).

These results offer new insight into multiple aspects of type IA topoisomerase activity. The differences in gate dynamics between topo III and topo I suggest a possible mechanism for their different biochemical activities and roles in the cell^14,15^. Whereas topo I removes negative supercoils, topo III functions primarily as a decatenase^15^, an activity which requires passing duplex DNA through the break in a single strand. The fast dynamics of topo I could favor efficient relaxation of negatively supercoiled DNA, whereas the slower closing rate of topo III could facilitate capture of double stranded DNA. This hypothesis is supported by the gate stabilizing interaction between the decatenation loop^51^ of topo III and an acidic loop in domain II observed in our simulations. A similar role in promoting the open state has been proposed for the eukaryotic protein RMI1, which is required for the decatenation activity of topo IIIα^23,52^. Once DNA has entered the cavity, it may disrupt this interaction, leading to fast closing of the gate and potentially facilitating the processive bursts of relaxation activity observed by Terekhova et al^39^.

Because of their importance in genome maintenance and replication, topoisomerases are a potential target for both anti-bacterial and anti-cancer drugs^53-55^. Topo I is the only type IA topoisomerase present in mycobacteria. *Mycobacterium tuberculosis* topo I has been validated genetically and chemically as a TB drug target ^56^-^58^ Poison inhibitors targeting type IA topoisomerases in other bacterial pathogens are also expected to be bactericidal since trapping of a small number of the covalent cleavage complexes on the chromosomal DNA would be sufficient for bacterial cell killing^53,59,60^. It would be highly desirable to identify poisons that can inhibit DNA ligation following the DNA cleavage step. Currently no drugs have been identified which target type IA topoisomerases. Design of such drugs has been hampered by the inability to experimentally probe the details of the full strand passage reaction. Key residues that drive the conformational changes for the opening and closing of the DNA "gate” have not yet been identified. The magnetic tweezers assays we developed reveal previously unmeasurable aspects of type IA topoisomerase activity. These assays have the potential to be powerful tools in testing the effects of mutations and small molecule interactions on this class of proteins.

## Methods

### Protein purification

Recombinant topoisomerase III was purified as previously described in Seol et al. (2013)^61^). Recombinant *E. coli* topoisomerase I and the Y319F mutant were purified as previously described^62^.

### DNA constructs for magnetic tweezers experiments

#### Hairpin construct

For the hairpin assay, we generated a 537-base hairpin with dsDNA handles on both ends as described in Seol et at. (2016)^63^.The top handle was modified with biotin on the 3’ end for attachment to a streptavidin coated magnetic bead and the bottom handle was modified with three digoxigenin at the 5' end for attachment to the coverslip. Oligonucleotides for hairpin generation were purchased from Eurofins MWG Operon.

#### Gapped DNA construct

Linear 2.5-kb dsDNA with a 37-nt gap with a single biotin or digoxigenin at each 5'-end was prepared as described in Seol et al. (2013)^61^. The2.5-kb PCR product was digested with Nt.BbvCI (New England Biolabs) to create two nicks at positions 2258 and 2294, near the middle of the DNA template. The ssDNA was engineered to have a CTTC sequence, which is expected to be preferentially cleaved by topoisomerases.

### Magnetic tweezers experiments

#### Magnetic tweezers sample cell preparation

Sample cell preparation is described in detail in Seol et al. (2016)^63^. Briefly, 5-10 × 10^−15^ mol DNA substrate was mixed with ∼100 ng anti-digoxygenin antibody (Sigma-Aldrich) and incubated in a sample cell overnight at 4° C. Unbound DNA was washed out with 200 μL of wash buffer (PBS supplemented with 0.04% v/v Tween-20 and 0.3% w/v BSA). 1-μm streptavidin-coated magnetic beads (MyOne, Invitrogen) were introduced into the sample cell and allowed to bind DNA for 5 minutes. The sample cell was then washed with 1 mL wash buffer, followed by 400 μL reaction buffer (50 mM Tris-HCL (pH 8.0), 30 mM monopotassium L-glutamic acid, 0.1 % Tween, 0.03% BSA, + an appropriate concentration of mM Mg L-glutamic acid). Measurements were taken at room temperature on a commercial Picotwist instrument. Bead position in three dimensions was acquired at 60 Hz.

#### Hairpin assay

Topo III or topo I was diluted to 1 μM in 0 Mg reaction buffer (50 mM Tris-HCL (pH 8.0), 30 mM monopotassium L-glutamic acid, 0.1 % Tween, 0.03% BSA) and incubated briefly at 37 °C. Samples were then diluted to 500 pM in 1 mM Mg reaction buffer. Hairpin DNA tethers were characterized before the addition of protein by applying a range of forces to determine the critical melting force of each individual tether, as slight differences in bead size could affect this value. 200 μL of was sample added to the experimental sample cell. A force of 8-10 pN was applied to determine a baseline for the folded state of the hairpin, and then a force or 20-24 pN was applied to manually melt the DNA. The DNA was held at high force for 30s - 5 min and then the force was returned to the baseline level. Upon full refolding of the hairpin, the force was again raised and the measurement repeated. Binding events were defined as pauses in refolding lasting > 100 ms.

#### Gapped DNA assay

Topoisomerase was diluted to 1 μM in 0 Mg reaction buffer and incubated briefly at 37 °C, then diluted to 250 pM in reaction buffer containing Mg L-glutamic acid. Forces of 8, 10, 12, 14, 16, and 18 pN were applied. Below 12 pN, closing rates for topo I were too fast to be sampled accurately with the time resolution of our experiments. Closing rates were too slow to provide adequate statistics at forces higher than 16 pN for topo III and 18 pN for topo I. Data was taken at 3 mM Mg^2+^ for topo III and 0.5 mM and 3 mM for topo I. For magnesium titration experiments, measurements were conducted at 8 pN on 250 pM topo III in reaction buffer containing 1 mM, 3 mM, 5 mM, or 10 mM Mg L-glutamic acid and at 14 pN for topo I at 0, 0.1 mM, 0.3 mM, 0.5 mM, and 3 mM Mg L-glutamic acid.

#### Molecular dynamics simulations

Simulations were conducted in NAMD^64^ v. 2.12 using the Charmm 36 force field^65,66^. Topo III bound to ssDNA (pdbid: I17D)^49^ was solvated with 148,705 TIP3P^67^ water molecules. Na and Cl ions were added to a concentration of 50 mM. The fully solvated system was minimized for 2500 steps with the protein atoms restrained with a force constant of 5 kcal/mol/Å^2^ and an additional 5000 steps without restraints. After minimization, the system was equilibrated at constant pressure and temperature for 500 ps. For the CPT equilibration step, pressure was held at 1.01325 Bar using Berendsen’s method and the protein atoms were harmonically restrained with a force constant of 5 kcal/mol/Å^2^. The full system was further equilibrated without restraints for 10 ns at constant volume and temperature. A Langevin thermostat was used to maintain a temperature of 300 K. From the closed structure, the protein was gradually opened by applying a harmonic restraint of 4 - 10 kcal/mol/Å^2^ on the distance between the center of mass of residues 290 to 415 and 492 to 620. The distance was gradually increased over a series of windows in 1 Å (0.1 nm) increments. Umbrella sampling windows were run for 2 ns - 20 ns, with a 200 ps equilibration period for each new window.

The reweighted free energy profile of the distance between the catalytic tyrosine and the cleavage site was calculated using a modification of the weighted histogram analysis method (WHAM)^48^ that allows for calculation of the free energy of a reaction coordinates orthogonal to the restrained coordinates^68^. Free Energy values were calculated with the program: WHAM: The weighted histogram analysis method, version 2.0.9.1, http://membrane.urmc.rochester.edu/content/wham^68^.

#### Data analysis

For magnetic tweezers hairpin experiments, data analysis was performed using Igor Pro 6.3A. For magnetic tweezers gapped experiments, traces were analyzed using vbFRET^38^ to assign states and then further analyzed in Igor Pro. Default values of priors were used in vbFRET analysis (upi=1, mu=0.5, beta=0.25, W=50, v=5, ua=1, uad=0). Lifetime analysis of the idealized traces from vbFRET were calculated using a thresholding algorithm. For molecular dynamics simulations, structural analysis was done using VMD. Free energy profiles were calculated using implementation of the weighted histogram analysis method, WHAM^68^. All errors are standard error unless otherwise noted.

## Data Availability

Source data for **Figure 4** is presented in **Supplementary Figures 3** and **4**. The data that support the findings of this study are available from the corresponding author upon reasonable request.

## Acknowledgments

This work was supported by the Intramural Research Program of the National Heart, Lung, and Blood Institute, National Institutes of Health (HL001056-07 to K.C.N.) and a grant from National Institutes of Health (R01GM054226 to Y.T).

## Author Contributions

M.M. and K.C. conceived the experiments. M.M. conducted the experiments and simulations and analyzed the data. Y.T.D. provided materials. M.M., K.C., and Y.T.D. wrote the manuscript.

## References

1. Champoux, J. J. DNA topoisomerases: structure, function, and mechanism. Annu. Rev. Biochem. 70, 369–413 (2001).

2. Corbett, K. D. & Berger, J. M. Structure, Molecular Mechanisms, and Evolutionary Relationships in DNA Topoisomerases. Annu. Rev. Biophys. Biomol. Struct. 33, 95–118 (2004).

3. Liu, L. F., Liu, C.-C. & Alberts, B. M. Type II DNA topoisomerases: Enzymes that can unknot a topologically knotted DNA molecule via a reversible double-strand break. Cell 19, 697–707 (1980).

4. Seol, Y., Zhang, H., Pommier, Y. & Neuman, K. C. A kinetic clutch governs religation by type IB topoisomerases and determines camptothecin sensitivity. Proc. Natl. Acad. Sci. 109, 16125–16130 (2012).

5. Stewart, L., Redinbo, M. R., Qiu, X., Hol, W. G. & Champoux, J. J. A model for the mechanism of human topoisomerase I. Science 279, 1534–1541 (1998).

6. Li, Z., Mondragón, A. & DiGate, R. J. The Mechanism of Type IA Topoisomerase-Mediated DNA Topological Transformations. Mol. Cell 7, 301–307 (2001).

7. Dekker, N. H. et al. The mechanism of type IA topoisomerases. Proc. Natl. Acad. Sci. 99, 12126–12131 (2002).

8. Tse-Dinh, Y. C. Biochemistry of bacterial type I DNA topoisomerases. Adv. Pharmacol. San Diego Calif 29A, 21–37 (1994).

9. Srivenugopal, K. S., Lockshon, D. & Morris, D. R. Escherichia coli DNA topoisomerase III: purification and characterization of a new type I enzyme. Biochemistry (Mosc.) 23, 1899–1906 (1984).

10. Confalonieri, F. et al. Reverse gyrase: a helicase-like domain and a type I topoisomerase in the same polypeptide. Proc. Natl. Acad. Sci. 90, 4753–4757 (1993).

11. Goulaouic, H. et al. Purification and characterization of human DNA topoisomerase IIIα. Nucleic Acids Res. 27, 2443–50 (1999).

12. Li, W. & Wang, J. C. Mammalian DNA topoisomerase IIIα is essential in early embryogenesis. Proc. Natl. Acad. Sci. 95, 1010–1013 (1998).

13. Seki, T., Seki, M., Onodera, R., Katada, T. & Enomoto, T. Cloning of cDNA Encoding a Novel Mouse DNA Topoisomerase III (Topo IIIβ) Possessing Negatively Supercoiled DNA Relaxing Activity, Whose Message Is Highly Expressed in the Testis. J. Biol. Chem. 273, 28553–28556 (1998).

14. Zechiedrich, E. L. et al. Roles of Topoisomerases in Maintaining Steady-state DNA Supercoiling in Escherichia coli. J. Biol. Chem. 275, 8103–8113 (2000).

15. Nurse, P., Levine, C., Hassing, H. & Marians, K. J. Topoisomerase III Can Serve as the Cellular Decatenase in Escherichia coli. J. Biol. Chem. 278, 8653–8660 (2003).

16. DiGate, R. J. & Marians, K. J. Identification of a potent decatenating enzyme from Escherichia coli. J. Biol. Chem. 263, 13366–13373 (1988).

17. Dean, F. B., Stasiak, A., Koller, T. & Cozzarelli, N. R. Duplex DNA knots produced by Escherichia coli topoisomerase I. Structure and requirements for formation. J. Biol. Chem. 260, 4975–4983 (1985).

18. Massé, E. & Drolet, M. Relaxation of transcription-induced negative supercoiling is an essential function of Escherichia coli DNA topoisomerase I. J. Biol. Chem. 274, 16654–16658 (1999).

19. Suski, C. & Marians, K. J. Resolution of Converging Replication Forks by RecQ and Topoisomerase III. Mol. Cell 30, 779–789 (2008).

20. Perez-Cheeks, B. A., Lee, C., Hayama, R. & Marians, K. J. A role for topoisomerase III in Escherichia coli chromosome segregation. Mol. Microbiol. 86, 1007–1022 (2012).

21. Cortés, F., Pastor, N., Mateos, S. & Domínguez, I. Roles of DNA topoisomerases in chromosome segregation and mitosis. Mutat. Res. Mutat. Res. 543, 59–66 (2003).

22. Goodwin, A., Wang, S.-W., Toda, T., Norbury, C. & Hickson, I. D. Topoisomerase III is essential for accurate nuclear division in Schizosaccharomyces pombe. Nucleic Acids Res. 27, 4050–4058 (1999).

23. Bocquet, N. et al. Structural and mechanistic insight into Holliday junction dissolution by Topoisomerase IIIα and RMI1. Nat. Struct. Mol. Biol. 21, 261–268 (2014).

24. Bussen, W., Raynard, S., Busygina, V., Singh, A. K. & Sung, P. Holliday junction processing activity of the BLM-Topo IIIαlpha-BLAP75 complex. J. Biol. Chem. 282, 31484–31492 (2007).

25. Ahmad, M. et al. Topoisomerase 3ß is the major topoisomerase for mRNAs and linked to neurodevelopment and mental dysfunction. Nucleic Acids Res. 45, 2704–2713 (2017).

26. Stoll, G. et al. Deletion of TOP3β, a component of FMRP-containing mRNPs, contributes to neurodevelopmental disorders. Nat. Neurosci. 16, 1228–1237 (2013).

27. Xu, D. et al. Top3β is an RNA topoisomerase that works with fragile X syndrome protein to promote synapse formation. Nat. Neurosci. 16, 1238–1247 (2013).

28. Viard, T. & de la Tour, C. B. Type IA topoisomerases: A simple puzzle? Biochimie 89, 456–467 (2007).

29. Lima, C. D., Wang, J. C. & Mondragón, A. Three-dimensional structure of the 67K N-terminal fragment of E. coli DNA topoisomerase I. Nature 367, 138–146 (1994).

30. Mondragón, A. & DiGate, R. The structure of Escherichia coli DNA topoisomerase III. Structure 7, 1373–1383 (1999).

31. Feinberg, H., Lima, C. D. & Mondragón, A. Conformational changes in E. coli DNA topoisomerase I. Nat. Struct. Biol. 6, 918–922 (1999).

32. Baker, N. M., Rajan, R. & Mondragón, A. Structural studies of type I topoisomerases. Nucleic Acids Res. 37, 693–701 (2009).

33. Leelaram, M. N. et al. Type IA topoisomerase inhibition by clamp closure. FASEB J. 27, 3030–3038 (2013).

34. Gunn, K. H., Marko, J. F. & Mondragón, A. An orthogonal single-molecule experiment reveals multiple-attempt dynamics of type IA topoisomerases. Nat. Struct. Mol. Biol. 24, 484–490 (2017).

35. Seol, Y. & Neuman, K. Magnetic Tweezers for Single-Molecule Manipulation. in Single Molecule Analysis (eds. Peterman, E. J. G. & Wuite, G. J. L.) 265–293 (Humana Press, 2011).

36. Mills, M. et al. RecQ helicase triggers a binding mode change in the SSB-DNA complex to efficiently initiate DNA unwinding. Nucleic Acids Res. 45, 11878–11890 (2017).

37. Perry, K. & Mondragón, A. Structure of a Complex between E. coli DNA Topoisomerase I and Single-Stranded DNA. Structure 11, 1349–1358 (2003).

38. Bronson, J. E., Fei, J., Hofman, J. M., Gonzalez, R. L. & Wiggins, C. H. Learning Rates and States from Biophysical Time Series: A Bayesian Approach to Model Selection and Single-Molecule FRET Data. Biophys. J. 97, 3196–3205 (2009).

39. Terekhova, K., Gunn, K. H., Marko, J. F. & Mondragón, A. Bacterial topoisomerase I and topoisomerase III relax supercoiled DNA via distinct pathways. Nucleic Acids Res. 40, 10432–10440 (2012).

40. Terekhova, K., Marko, J. F. & Mondragón, A. Single-molecule analysis uncovers the difference between the kinetics of DNA decatenation by bacterial topoisomerases I and III. Nucleic Acids Res. gku785 (2014).

41. Zhang, J., Pan, B., Li, Z., Sheng Zhao, X. & Huang, L. Kinetic insights into the temperature dependence of DNA strand cleavage and religation by topoisomerase III from the hyperthermophile Sulfolobus solfataricus. Sci. Rep. 7, 5494 (2017).

42. Domanico, P. L. & Tse-Dinh, Y. C. Mechanistic studies on E. coli DNA topoisomerase I: Divalent ion effects. J. Inorg. Biochem. 42, 87–96 (1991).

43. Tse-Dinh, Y. C. Uncoupling of the DNA breaking and rejoining steps of Escherichia coli type I DNA topoisomerase. Demonstration of an active covalent protein-DNA complex. J. Biol. Chem. 261, 10931–10935 (1986).

44. Sorokin, E. P. et al. Inhibition of Mg2+ binding and DNA religation by bacterial topoisomerase I via introduction of an additional positive charge into the active site region. Nucleic Acids Res. 36, 4788–4796 (2008).

45. Zhu, C. X., Roche, C. J. & Tse-Dinh, Y. C. Effect of Mg(II) binding on the structure and activity of Escherichia coli DNA topoisomerase I. J. Biol. Chem. 272, 16206–16210 (1997).

46. Bhat, A. G., Leelaram, M. N., Hegde, S. M. & Nagaraja, V. Deciphering the distinct role for the metal coordination motif in the catalytic activity of Mycobacterium smegmatis topoisomerase I. J. Mol. Biol. 393, 788–802 (2009).

47. Torrie, G. M. & Valleau, J. P. Nonphysical sampling distributions in Monte Carlo free-energy estimation: Umbrella sampling. J. Comput. Phys. 23, 187–199 (1977).

48. Souaille, M. & Roux, B. Extension to the weighted histogram analysis method: combining umbrella sampling with free energy calculations. Comput. Phys. Commun. 135, 40–57 (2001).

49. Changela, A., DiGate, R. J. & Mondragón, A. Crystal structure of a complex of a type IA DNA topoisomerase with a single-stranded DNA molecule. Nature 411, 1077–1081 (2001).

50. Minh, D. D. L. Multiple Potentials of Mean Force from Biased Experiments Along a Single Coordinate. J. Phys. Chem. B 111, 4137–4140 (2007).

51. Li, Z., Mondragón, A., Hiasa, H., Marians, K. J. & DiGate, R. J. Identification of a unique domain essential for Escherichia coli DNA topoisomerase III-catalysed decatenation of replication intermediates. Mol. Microbiol. 35, 888–895 (2000).

52. Cejka, P., Plank, J. L., Dombrowski, C. C. & Kowalczykowski, S. C. Decatenation of DNA by the S. cerevisiae Sgs1-Top3-Rmi1 and RPA complex: a mechanism for disentangling chromosomes. Mol. Cell 47, 886–896 (2012).

53. Tse-Dinh, Y.-C. Targeting bacterial topoisomerase I to meet the challenge of finding new antibiotics. Future Med. Chem. 7, 459–471 (2015).

54. Sandhaus, S. et al. Small-Molecule Inhibitors Targeting Topoisomerase I as Novel Antituberculosis Agents. Antimicrob. Agents Chemother. 60, 4028–4036 (2016).

55. Giles, G. I. & Sharma, R. P. Topoisomerase enzymes as therapeutic targets for cancer chemotherapy. Med. Chem. Shariqah United Arab Emir. 1, 383–394 (2005).

56. Nagaraja, V., Godbole, A. A., Henderson, S. R. & Maxwell, A. DNA topoisomerase I and DNA gyrase as targets for TB therapy. Drug Discov. Today 22, 510–518 (2017).

57. Ahmed, W., Menon, S., Godbole, A. A., Karthik, P. V. D. N. B. & Nagaraja, V. Conditional silencing of topoisomerase I gene of Mycobacterium tuberculosis validates its essentiality for cell survival. FEMS Microbiol. Lett. 353, 116–123 (2014).

58. Ravishankar, S. et al. Genetic and chemical validation identifies Mycobacterium tuberculosis topoisomerase I as an attractive anti-tubercular target. Tuberc. Edinb. Scotl. 95, 589–598 (2015).

59. Aedo, S. & Tse-Dinh, Y.-C. Isolation and Quantitation of Topoisomerase Complexes Accumulated on Escherichia coli Chromosomal DNA. Antimicrob. Agents Chemother. 56, 5458–5464 (2012).

60. Pommier, Y. & Marchand, C. Interfacial inhibitors: targeting macromolecular complexes. Nat. Rev. Drug Discov. 11, 25–36 (2011).

61. Seol, Y., Hardin, A. H., Strub, M.-P., Charvin, G. & Neuman, K. C. Comparison of DNA decatenation by Escherichia coli topoisomerase IV and topoisomerase III: implications for non-equilibrium topology simplification. Nucleic Acids Res. gkt136 (2013).

62. Zhu, C.-X. & Tse-Dinh, Y.-C. Overexpression and Purification of Bacterial DNA Topoisomerase I. in DNA Topoisomerase Protocols 145–151 (Humana Press, 1999).

63. Seol, Y., Strub, M.-P. & Neuman, K. C. Single molecule measurements of DNA helicase activity with magnetic tweezers and t-test based step-finding analysis. Methods San Diego Calif 105, 119–127 (2016).

64. Kalé, L. et al. NAMD2: Greater Scalability for Parallel Molecular Dynamics. J. Comput. Phys. 151, 283–312 (1999).

65. Brooks, B. R. et al. CHARMM: The Biomolecular Simulation Program. J. Comput. Chem. 30, 1545–1614 (2009).

66. Huang Jing & MacKerell Alexander D. CHARMM36 all-atom additive protein force field: Validation based on comparison to NMR data. J. Comput. Chem. 34, 2135–2145 (2013).

67. Neria, E., Fischer, S. & Karplus, M. Simulation of activation free energies in molecular systems. J. Chem. Phys. 105, 1902–1921 (1996).

68. Grossfield, A. WHAM: The weighted histogram analysis method, version 2.0.9.1, http://membrane.urmc.rochester.edu/content/wham.

